# Short-term Effects of Vagus Nerve Stimulation on Learning and Evoked Activity in Auditory Cortex

**DOI:** 10.1101/2019.12.19.883165

**Authors:** Jesyin Lai, Stephen V. David

## Abstract

Chronic vagus nerve stimulation (VNS) can facilitate learning of sensory and motor behaviors. VNS is believed to trigger release of neuromodulators, including norepinephrine and acetylcholine, which can mediate cortical plasticity associated with learning. Most previous work has studied effects of VNS over many days, and less is known about how acute VNS influences neural coding and behavior over the shorter term. To explore this question, we measured effects of VNS on learning of an auditory discrimination over 1-2 days. Ferrets implanted with cuff electrodes on the vagus nerve were trained by classical conditioning on a tone frequency-reward association. One tone was associated with reward while another tone, was not. The frequencies and reward associations of the tones were changed every two days, requiring learning of a new relationship. When the tones (both rewarded and non-rewarded) were paired with VNS, rates of learning increased on the first day following a change in reward association. To examine VNS effects on auditory coding, we recorded single- and multi-unit neural activity in primary auditory cortex (A1) of passively listening animals following brief periods of VNS (20 trials/session) paired with tones. Because afferent VNS induces changes in pupil size associated with fluctuations in neuromodulation, we also measured pupil during recordings. After pairing VNS with a neuron’s best-frequency (BF) tone, responses in a subpopulation of neurons were reduced. Pairing with an off-BF tone or performing VNS during the inter-trial interval had no effect on responses. We separated the change in A1 activity into two components, one that could be predicted by fluctuations in pupil and one that persisted after VNS and was not accounted for by pupil. The BF-specific reduction in neural responses remained, even after regressing out changes that could be explained by pupil. In addition, the size of VNS-mediated changes in pupil predicted the magnitude of persistent changes in the neural response. This interaction suggests that changes in neuromodulation associated with arousal gate the long-term effects of VNS on neural activity. Taken together, these results support a role for VNS in auditory learning and help establish VNS as a tool to facilitate neural plasticity.

## 1 INTRODUCTION

The vagus nerve (cranial nerve X) is a major component of the autonomic nervous system. In addition to modulating activity in the heart, lungs and digestive tract via efferent fibers, the vagus nerve also conveys signals from the head, neck, and body to the central nervous system via afferent fibers, which constitute 80 % of the nerve (Foley and DuBois, 1937). Stimulation of these afferent fibers has been used in clinical therapies for epilepsy, depression and other neurological disorders (Groves and Brown, 2005). Afferent signals activate widespread release of norepinephrine and acetylcholine in the brain via the nucleus tractus solitarius, the locus coeruleus (LC) and nucleus basalis (NB) (Engineer et al., 2013; Dorr and Debonnel, 2006). Increased release of these neuromodulators is associated with heightened arousal and has an impact in modulating learning and memory (McGaugh, 1989). Chronic vagus nerve stimulation (VNS) has been reported to improve learning and memory of associated events in humans (Clark et al., 1999) and rats (Clark et al., 1995).

Previous studies have demonstrated that VNS during rehabilitative training improves recovery of motor function in several models of brain injury (Khodaparast et al., 2016, 2014; Hays et al., 2014a,b, 2016; Meyers et al., 2018; Morrison et al., 2019). The therapeutic effects of pairing VNS with motor rehabilitation persist even after the cessation of stimulation, suggesting that the VNS-induced plasticity and learning are long-term (Khodaparast et al., 2014; Hays et al., 2014a). VNS has also been reported to enhance memory and to facilitate extinction of fear conditioning in rats (Pena et al., 2013, 2014). These diverse findings suggest that VNS supports a general facilitation of skill learning, which could also include learning of new auditory categories. Recruitment of neuromodulatory activity by VNS is believed to contribute to this enhanced learning (Engineer et al., 2015a). However, the number of studies investigating VNS effects on auditory learning and the mechanisms of VNS is limited.

Activation of neuromodulatory systems has been broadly implicated in neural plasticity. For example, acute stimulation of NB causes widespread release of acetlycholine and can enhance the reliability of sensory coding (Goard and Dan, 2009) as well as promote learning and memory (Miasnikov et al., 2008). Repeatedly pairing auditory stimuli with electrical stimulation of NB in animals can produce shifts in auditory tuning that enhance cortical responses to paired stimuli, with large-scale, persistent changes in cortical receptive field organization (Kilgard and Merzenich, 1998; Bakin and Weinberger, 1996; Froemke et al., 2007). Similarly, repeatedly pairing a tone with LC stimulation induces selective plasticity for the paired frequency in auditory cortex (A1) (Edeline et al., 2011; Glennon et al., 2019). However, NB or LC stimulation is usually performed via highly invasive deep brain stimulation, which could be harmful to the brain and thus has limited therapeutic potential. Because VNS may generate neural plasticity similar to that associated with NB stimulation (Hays et al., 2013), it may provide a less invasive means of stimulation that bypasses the need for deep brain stimulation.

Recent work has shown that some changes in neuromodulatory activity are reflected in luminance-independent changes in pupil size (Joshi et al., 2016; Murphy et al., 2014; Desbeaumes Jodoin et al., 2015). Fluctuations in pupil diameter have been shown to correlate with changes in sensory spiking activity (McGinley et al., 2015; Vinck et al., 2015) and with adrenergic and cholinergic axon terminal activity in cortex (Reimer et al., 2016). As pupil dilation is observed following VNS (Bianca and Komisaruk, 2007; Desbeaumes Jodoin et al., 2015) and has been proposed as a readout of VNS efficacy (Mridha et al., 2019), we speculate that changes in the activity of A1 neurons following pairing of VNS with an acoustic stimulus could be mediated by two pathways (Fig. 1). First, short-term, reversible changes in excitability should correlate with pupil diameter (Schwartz et al., 2019). Second, long-term, persistent changes in neural activity should reflect synaptic plasticity and learning, and should last longer than changes in pupil. These pathways may not be independent, as the long-term effects may be gated by the short-term fluctuations in neuromodulatory activity.

**Figure 1.**
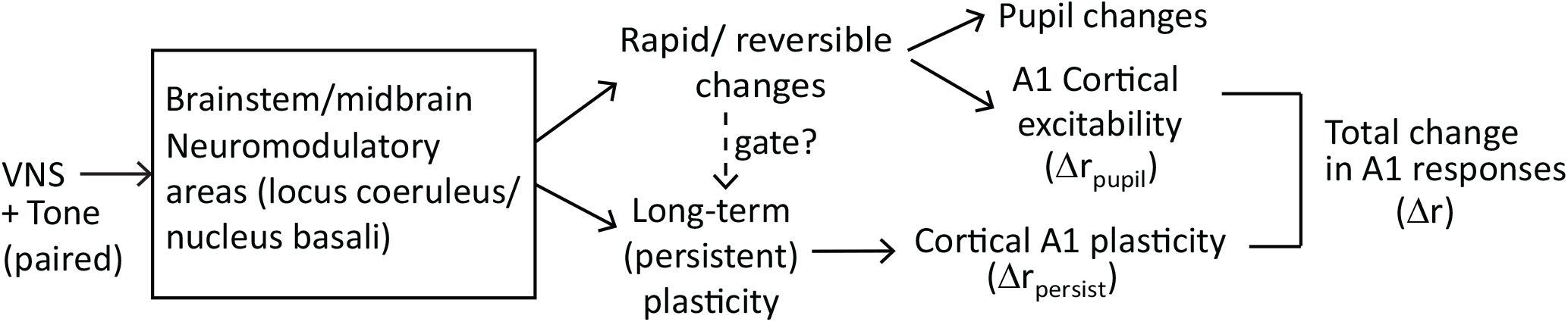
Two possible pathways by which VNS could mediate changes in A1 spiking activity. In the short-term, reversible pathway, VNS evokes changes in pupil that are correlated with neural excitability. In the long-term, persistent pathway, which maybe gated by the rapid and reversible changes, VNS produces long-term plasticity related to learning. The net change in the A1 response is the sum of short-term and long-term effects, Δ*r* = Δ*r*_*pupil*_ + Δ*r*_*persist*_.

Although previous studies have shown plasticity in A1 following VNS, most performed chronic pairing of VNS with sound presentation over many days, e.g., 300 times/day for 20 days Engineer et al. (2015b); Shetake et al. (2012); Engineer et al. (2017, 2011); Borland et al. (2019). Here, we studied short-term effects of VNS on auditory learning and cortical plasticity in A1. To study behavioral effects of VNS, we developed a paradigm for reward association learning over 200-500 trials (1-2 days). Ferrets were trained by classical conditioning to discriminate between rewarded and non-rewarded tones, and we compared the rate of learning with and without VNS. Learning rates increased when VNS was paired with task stimuli but not when VNS occurred during inter-trial intervals.

To measure effects of VNS on cortical activity, we recorded single- and multi-unit neural activity in A1 of passively listening animals. VNS was paired with tone stimuli similar to those used for behavior. Neural responses in a subpopulation of neurons were decreased after pairing VNS with the best frequency (BF) tone. To determine the interaction between VNS, pupillometry and neurophysiological activity, we recorded pupil during the passive stimulation experiments. The pupil data revealed that trial-to-trial variability in neural responses was correlated with pupil size. Regressing out effects of pupil-indexed arousal decreased the difference in spiking activity of post- versus pre-VNS. However, neural responses of post-VNS remained significantly reduced. Together, these results demonstrate a role of VNS in facilitating learning and promoting neural plasticity in auditory cortex.

## 2 METHODS AND MATERIALS

### 2.1 Ethics statement

All procedures were approved by the Oregon Health and Science University Institutional Animal Care and Use Committee (protocol IP00001561) and conform to standards of the Association for Assessment and Accreditation of Laboratory Animal Care (AAALAC).

### 2.2 Animals

Three young adult male ferrets (animals P, S, and N) were obtained from an animal supplier for use in this study (Marshall Farms, New York). Prior to experiments, animals were implanted surgically with a head post for head fixation and to expose a portion of the skull for access to auditory cortex. Anesthesia was induced using Ketamine (35 mg/kg, intramascular (IM) injection) and Xylazine (5 mg/kg IM) and maintained with isoflurane (0.5-2 %). A warmed saline solution (10 mL) was given to the animals to prevent dehydration. Anesthesia depth was monitored by heart rate, respiration rate and blood oxygen percentage. Under sterile conditions, the head post was mounted to the skull using dental acrylic (AM Systems) or Charisma composite, which bonded to the skull and to a set of stainless steel screws embedded in the bone. After the surgery, animals were treated with prophylactic antibiotics (Baytril at 100 mg/ml, subcutaneous (SC) injection) and analgesics under the supervision of University veterinary staff. The wound was cleaned and bandaged during a two-week recovery period. After recovery, each ferret was gradually acclimated to head fixation using a custom stereotaxic apparatus in a plexiglass tube. Habituation sessions initially lasted for 5 minutes and increased by increments of 5–10 minutes daily until the ferret lay comfortably in the tube for at least one hour.

### 2.3 Vagus nerve implant surgery

After acclimation to head fixation, each animal was implanted with a tripolar cuff electrode (Cortec or Microprobes) around the left cervical vagus nerve. Induction and other aspects of the surgery were the same as for the head post implant.

Animals were placed in a supine position for implantation of the electrode cuff. Lidocaine (2 mg/kg SC) was injected in the neck at the incision site, and the left cervical vagus nerve was exposed through blunt dissection of the neck. The cuff was secured around the nerve (Surgical protocol adapted from a VNS study in rats (Lu et al., 2018)). Leads from the electrode were tunneled subcutaneously from the implant site in the neck to the head post. The leads for the cuff electrode were secured to the head post implant with acrylic.

After connecting the cuff pins to an electrical stimulator (AM systems 2100), efficacy of VNS was confirmed by observation of a heart rate drop while stimulating the nerve. Upon confirmation that the cuff was successfully stimulating the nerve, the neck was sutured closed and Bacitracin antibiotic cream was applied to incision sites on the neck and head. Animals were given Baytril (10 mg/kg SC) and Buprenorphine (0.02 mg/kg SC) for 2 days after surgery.

### 2.4 Validation of cuff electrode function

To confirm the function of the implanted cuff following implantation, we performed VNS and simultaneously measured heart rate decreases (under anesthesia) or pupil size changes (awake, passive conditions). For heart rate measurement, animals were anesthetized with Ketamine (5 mg/kg IM) and Dexmedetomidine (0.05 mg/kg IM). Atropine (0.05 mg/kg IM) was injected to prevent bradycardia. Animals’ heart rate was measured immediately before and after delivery of current to the cuff (0.1-2 mA, 200 *μ*s biphasic pulses at 30 Hz, 3-5 s duration). For pupil size measurement, animals were head-fixed and pupillometry was obtained by infrared video before and after VNS, as described below.

### 2.5 Pupillometry

During VNS and neurophysiological recordings, changes of pupil in one eye of animals were recorded as video using an open-source camera (Adafruit TTL Serial Camera) fitted with a lens (M12 Lenses PT-2514BMP, 25 mm). The camera was placed at approximately 10 cm from the animal’s eye. To improve contrast, the imaged eye was illuminated by a bank of infrared LED light. Constant light intensity was fixed and provided using a ring light (AmScope LED-144S) so that a maximum dynamic range of pupil sizes could be measured.

Pupil size was measured using custom written code in MATLAB or Python. The details of pupil diameter measurement in early recordings are similar to Schwartz *et al.* (2019). For the later recordings using Python analysis, we trained a machine learning algorithm to locate the pupil in each video frame and fit an ellipse to the pupil boundary. The algorithm was trained on video frames that were labeled using the methods in Schwartz *et al.* (2019). After training, the model performed well on novel video frames from new animals. If the fit quality was poor, the model was retrained and the analysis rerun to obtain a better fit quality. The code for this analysis is available at https://github.com/LBHB/nems_db.

For both analyses, we defined pupil size as the length of the minor axis (in pixels) of the fit ellipse. The frame rate of the cameras varied from 10 to 30 frames/second. A timestamp was recorded at the start and end of each trial with subsequent interpolated measurements of pupil size to match the sampling of the simultaneously recorded neural data. This procedure ensured that the two data streams (video and neural recording) remained synchronized throughout each recording.

To remove blink artifacts, rapid and transient changes in pupil size were identified (McGinley et al., 2015). The derivative of the pupil trace was taken and bins with derivatives more than 6 standard deviations from the mean were marked. Blinks were identified within these bins by screening for decreases in pupil size followed by increases. Data during a 6-second period surrounding the blink was then removed from the trace and replaced by a linear interpolation of the pupil size immediately before and after the blink. Pupil data was shifted by 750 ms relative to spike times in order to account for the lagged relationship between changes in pupil size and neural activity in auditory cortex to allow for comparison with previous research (McGinley et al., 2015).

### 2.6 Target-reward association task

Following recovery from cuff implant surgery, the animals were trained to learn a tone frequency-reward association using classical conditioning. Its behavioral data was not included in this study. On each behavioral trial, a pure tone target (T1, 1 s, 60 dB SPL) was presented at a random time in a sequence of broadband noise distractor sounds (temporally orthogonal ripple combinations, TORCS (Klein et al., 2000), 1 s duration, 1 s inter-stimulus interval). A reward (0.8-1.5 mL Ensure) was delivered immediately after target offset. Liquid reward delivery was controlled electronically with a solenoid valve. Trials proceeded independently of animal behavior, but licks were monitored by a piezo resistor attached to the lick spout. Licking after tone onset but before reward delivery indicated anticipation of the reward and were interpreted as learning of the reward association. When animals learned to associate T1 with a liquid reward, another tone (T2, frequency 2-3 octaves away from T1) was presented on randomly interleaved trials but with no reward. In classical condition protocols, T1 and T2 are often referred to as CS+ and CS-, respectively. The frequencies of T1 and T2 were changed every 2 days (200-250 trials/day), which is the average time animals took to discriminate the rewarded versus non-rewarded target without VNS pairing.

After animals demonstrated an ability to learn the target-reward associations, VNS was introduced during behavior. Two conditions were tested, tone-paired and -unpaired. In the paired condition, both T1 and T2 were paired with VNS (1 s duration, 30 Hz, 200 *μ*s biphasic pulses, 0.4-2 mA, VNS starting 100-150 ms before T1/T2 onset). In the VNS unpaired condition, VNS occurred randomly during the inter-trial interval silence after T1 or T2 presentation. Basic stimulation parameters were taken from Shetake *et al.* (2012). However, the stimulation current for VNS was set for each animal by adjusting the current from 0.4 mA to 2.0 mA in 0.2 mA steps. The lowest effective current was determined as the level at which the animal showed higher cumulative lick rate on the rewarded tone for paired versus unpaired VNS-tone condition. Once this level was determined for each animal, the same current was then used in all the training sessions and recordings for data acquisition (animal P: 1.5 mA, S: 2.0 mA, and N: 0.4 mA). The impedance of the cuff electrode (5-15 *k*Ω) was verified during training by converting the voltage required to produce the stimulation current into a resistance value. All data reported in the results were collected after the effective stimulation current was established.

To quantify animals’ learning to discriminate the rewarded and non-rewarded target tones, we calculated learning index (LI) with the below equation:

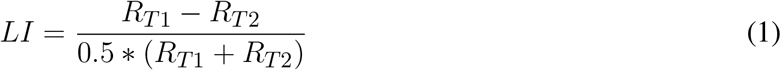

*R* is the probability of licking during the 0.95 s window from 0.2 s after tone onset to 0.15 s after tone offset. A value of *LI* > 0 indicates that the animal is preferentially responding to the rewarded tone, T1. No feedback was given in response to licks. Lick rates tended to be lower during the presentation of the noise distractors, but those data are not relevant to the learning analysis and are not reported here.

### 2.7 Neurophysiology

To allow for neurophysiological recordings, a microcraniotomy was opened over primary auditory cortex (A1). Extracellular neurophysiological activity was recorded using 1-2 tungsten microelectrodes (FHC) or a 64-channel electrode array (Masmanidis Lab, UCLA (Yang et al., 2019)). Both the microelectrodes and array were inserted into A1 with independent motorized microdrives (Alpha-Omega EPS). Amplified (AM Systems 3600) and digitized (National Instruments) signals were stored using open-source data acquisition software (Englitz et al., 2013). Recording sites were confirmed as being in primary auditory cortex based on tonotopy and relatively reliable and simple response properties (Shamma et al., 1993; Atiani et al., 2014). During the recording session, animals were observed and monitored by video camera. Acoustic stimulus presentation was controlled by custom MATLAB software (https://bitbucket.org/lbhb/baphy). Digital acoustic signals were transformed to analog (National Instruments), amplified (Crown),and delivered through a free-field speaker (Manger). Pupillometry was performed during neurophysiological recordings, as described above.

To isolate a spiking unit while positioning tungsten electrodes, a pure-tone or broadband noise probe stimulus was played periodically to search for sound-activated neurons. During recordings using 64-channel array, the probe was inserted into auditory cortex until neural activity was observed across the 1.05-mm span of recorded channels. A series of brief pure tone (100 ms duration at 60 dB SPL) was used to determine the best frequency (BF), *i.e.*, the frequency that evoked the strongest spike rate response.

After characterizing tuning properties at a recording site, two frequencies were selected for probing the effects of VNS on tone-evoked activity. One tone was fixed at BF and other 2-3 octaves away from BF (off-BF). The same tones were presented to passively listening animals in experimental blocks before, during and after a VNS session. The inter-trial interval was 12 seconds, each tone was presented 20 times per block. In the during-VNS block, VNS was either (1) paired with BF tone, (2) paired with off-BF tone or (3) unpaired with BF tone (VNS onset 6 s after tone onset). The electrophysiological amplifier was removed during the VNS session to prevent current leakage. Thus spiking data was not acquired during the VNS session.

Neurophysiological data was processed offline to identify spike events. For tungsten electrode recordings, the raw signals were band-pass filtered at 300–6000 Hz and then applying PCA-based clustering algorithm to spike-threshold events (David et al., 2009). For array recordings, single- and multi-units were sorted offline using Kilosort (Pachitariu et al., 2016). Neurons were considered isolated single units if standard deviation of spike amplitude was at least two times the noise floor, corresponding to >95 % isolation of spikes.

### 2.8 Evoked activity analysis

Spike rate and pupil data was binned at 30-50 Hz. Peri-stimulus time histogram (PSTH) responses to each tone were calculated by aligning spike activity to tone onset and averaging across repetitions. To obtain evoked activity, the mean spike rate during the 0.5 s silence (baseline spontaneous activity) preceding tone onset was subtracted from the PSTH.

Changes in the PSTH response to the BF tone (pre- versus post-VNS) were measured in each of the three VNS conditions. The PSTH was divided into 4 epochs: spontaneous activity (0-500 ms before tone onset), onset response (0-60 ms after tone onset), sustained response (60-1000 ms after tone onset) and offset response (0-100 ms after tone offset). Average spontaneous rate was subtracted from the other three responses. Changes in onset and offset responses were weak, and the results focus on changes in the sustained response.

### 2.9 Isolation of pupil-related changes by linear regression

Given our observation that pupil dilation is observed following VNS and changes in pupil correlate with changes in neural excitability in A1 (Schwartz et al., 2019), we sought to dissociate effects of VNS that could and could not be explained by changes in pupil. To accomplish this, we fitted a linear regression model using 40-fold validation to predict response due to pupil effects alone. This response (*r*_*pupil*_) was estimated using the recorded neural (*r*) and pupil (*p*) data.

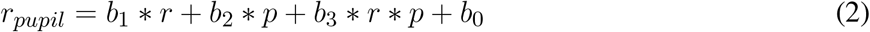

The change in response that persisted after accounting for pupil changes (*r*_*persist*_) was obtained by subtracting *r*_*pupil*_ from *r*. After regressing out the effects of pupil, a comparison of *r*_*persist*_ before and after VNS was performed similarly to the comparison of raw evoked responses before and after VNS described above.

## 3 RESULTS

Three ferrets (animals P, S, and N) were implanted with a cuff electrode for VNS. They were trained to perform the target-reward association task using classical conditioning (Fig. 2). Finally, single-unit neurophysiological data was also recorded from these animals.

**Figure 2.**
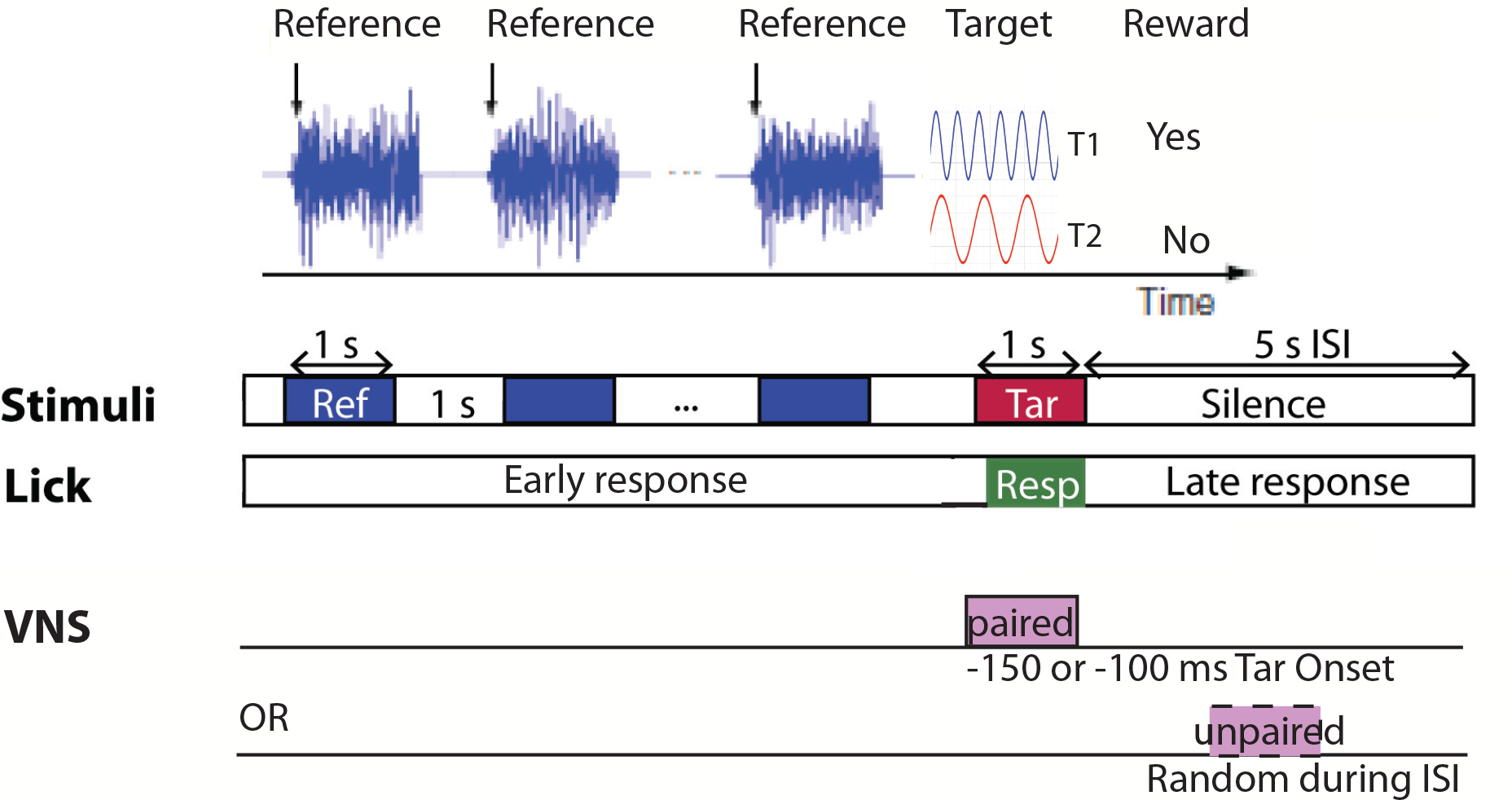
Animals were trained using classical conditioning to associate one specific target tone (T1) with a reward and another target tone (T2) with no reward. T1 and T2 were changed every 2 days (200-250 trials/day), typically after target-reward associations were learned. During these 2 days of training, T1 and T2 were either paired with VNS (1 s duration, 30 Hz, 200 *μ*s biphasic pulses, 0.4-2 mA, VNS onset 100-150 ms before T1/T2 onset) or unpaired with VNS (occurred randomly during inter-stimulus interval). Only lick(s) that occurred during target presentation (0.2-1.15 s after tone onset) were considered as a response to the target sound.

### 3.1 Pupil dilation following VNS

After animals recovered from the vagus nerve implant surgery, proper function of the cuff electrode was validated by measuring changes in heart rate and pupil size following VNS. Both heart rate reduction (Nearing et al., 2016) and pupil dilation (Desbeaumes Jodoin et al., 2015; Bianca and Komisaruk, 2007) in response to VNS have been reported previously. We measured pupil size fluctuations over time with and without 1 sec stimulation of the vagus nerve. We observed an increase in pupil diameter lasting for several seconds following VNS as compared to the control (pupil recorded during silence and without VNS, Fig. 3).

**Figure 3.**
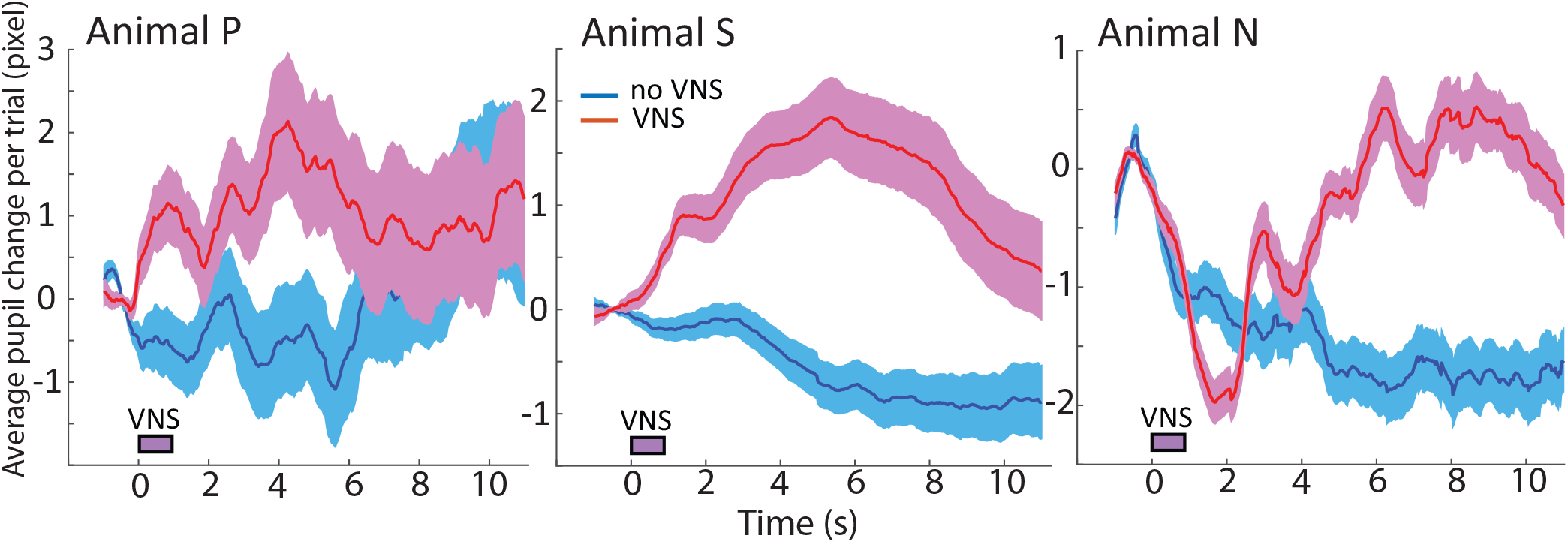
Pupil dilation followed VNS was observed in awake animals. Pupil size was recorded from animals under passive conditions with VNS but no sound presentation. Dilation time course and VNS threshold were different among animals. Shading indicate standard error of the mean.

### 3.2 Tone-VNS pairing improved target-reward association learning

We trained the animals to associate a tone T1 with a reward and T2 with no reward by classical conditioning. T1 and T2 frequencies and associated reward values were changed every 2 days (200-250 trials/day). An example plot of cumulative lick rate following target onset shows that animals could learn the reward categories correctly in 2 days under the control condition, *i.e.*, when VNS was not paired with targets but instead occurred during the inter-trial interval (Fig. 4A). In contrast, when VNS was paired with the presentation of the target tones (both T1 and T2), animals responded to T1 more than T2 on day 1 (trials 1-200, Fig. 4B). On day 2 (trials 201-400), the difference in cumulative response rates for T1 and T2 was even greater.

**Figure 4.**
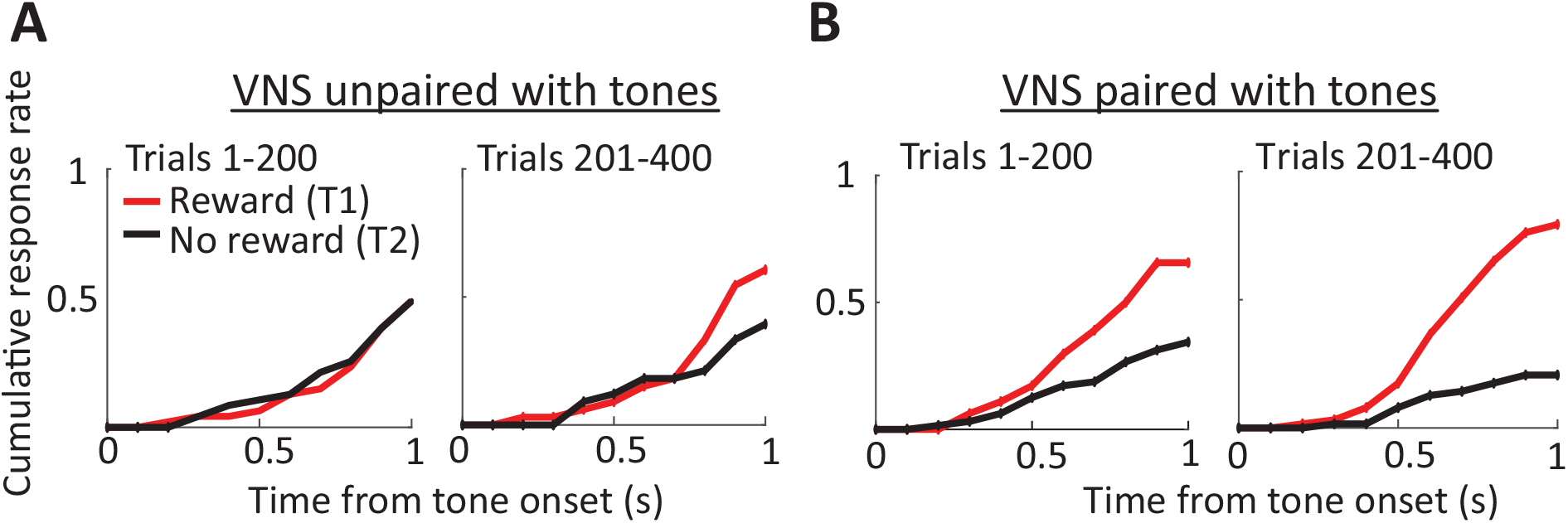
Training improved the speed of target reward association when VNS was paired with targets. Cumulative response rate as a function of time from target onset showed no difference between T1 and T2 on day 1 when VNS was unpaired with targets. Higher cumulative response rate was observed on day 2. In contrast, higher cumulative response rate was already noticed on day 1 when VNS was paired with targets.

Behavioral data for one animal across multiple reward pairing and VNS conditions are summarized in Fig. 5A. In addition to changing target frequencies pseudo-randomly every 2 days, we varied paired and unpaired VNS conditions as well in order to balance transition probabilities. Multiple training days were carried out to cover all the different combinations of target reward types and VNS conditions. Task conditions were changed every two days regardless of performance in order to prevent overtraining on a single target-reward association. Performance was measured using a learning index (LI, Eq. 1), computed as the normalized ratio of responses to T1 (rewarded) versus T2 (non-rewarded). An LI of 2 indicated responses only to T1, and a value of 0 indicated equal likelihood of response to T1 or T2. In training days when VNS was paired with targets, LI was mostly positive and higher than for unpaired VNS-tone sessions.

**Figure 5.**
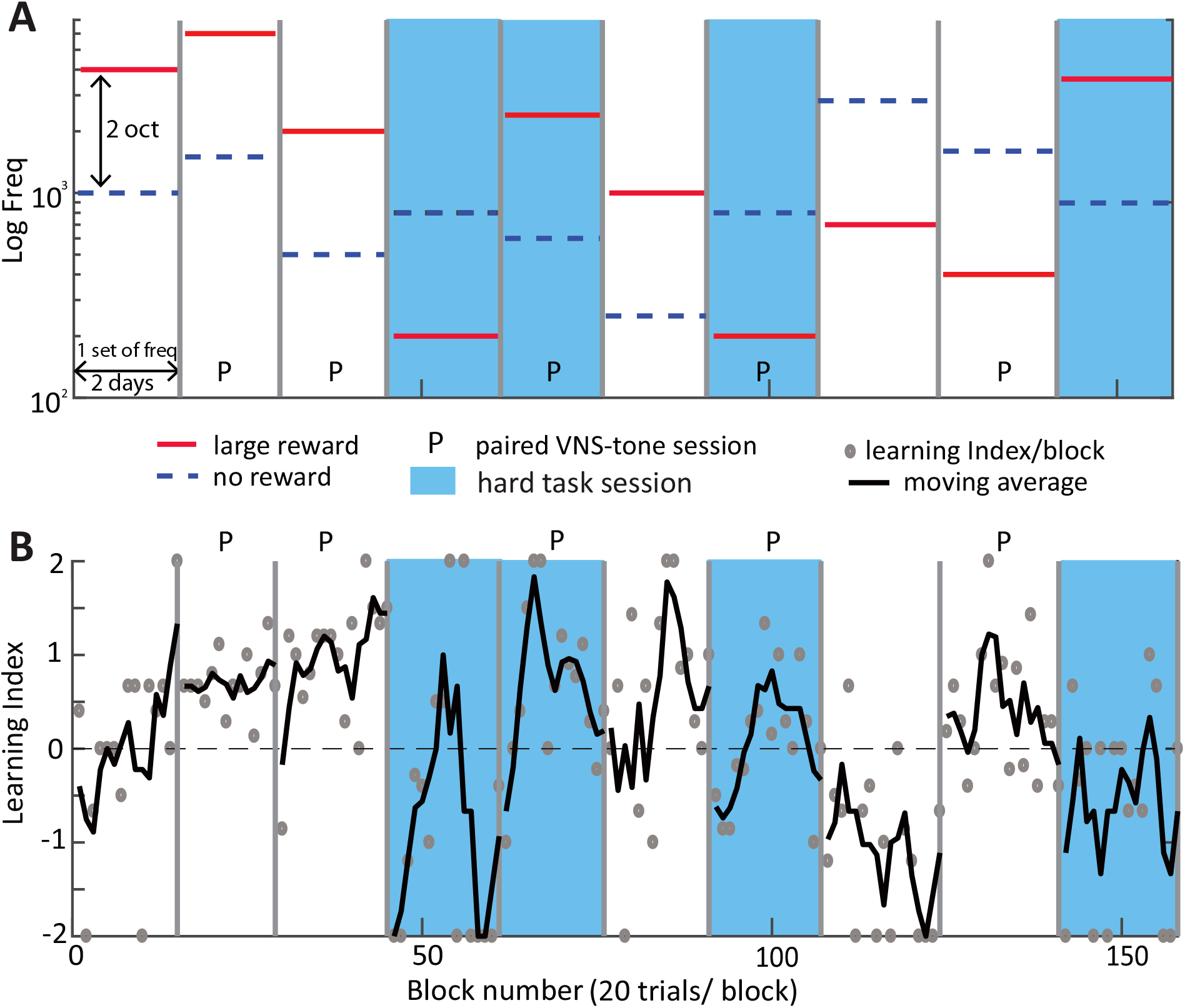
Learning index (LI) was mostly positive and higher for training days when VNS and tone presentation were paired (P) compared to when they were unpaired (N). P and N conditions were switched pseudo-randomly every 2 days so that they had balanced transition probability. Blue shading indicates training days when the task was considered hard because the rewarded tone (T1) was changed from the higher to the lower frequency than T2, or vice versa.

In addition to controlling for the VNS pairing condition, we also controlled for transitions between whether the lower- or higher frequency tone was rewarded. Even if absolute tone frequency changed, animals had a more difficult time learning reward associations if T1 switched from being higher frequency than T2 to lower frequency than T2, or vice versa (blue shading in Fig. 5A). Transitions in which the relative frequency of T1 to T2 switched were considered difficult and those in which it did not switch were considered easy (see Fig. 6B).

**Figure 6.**
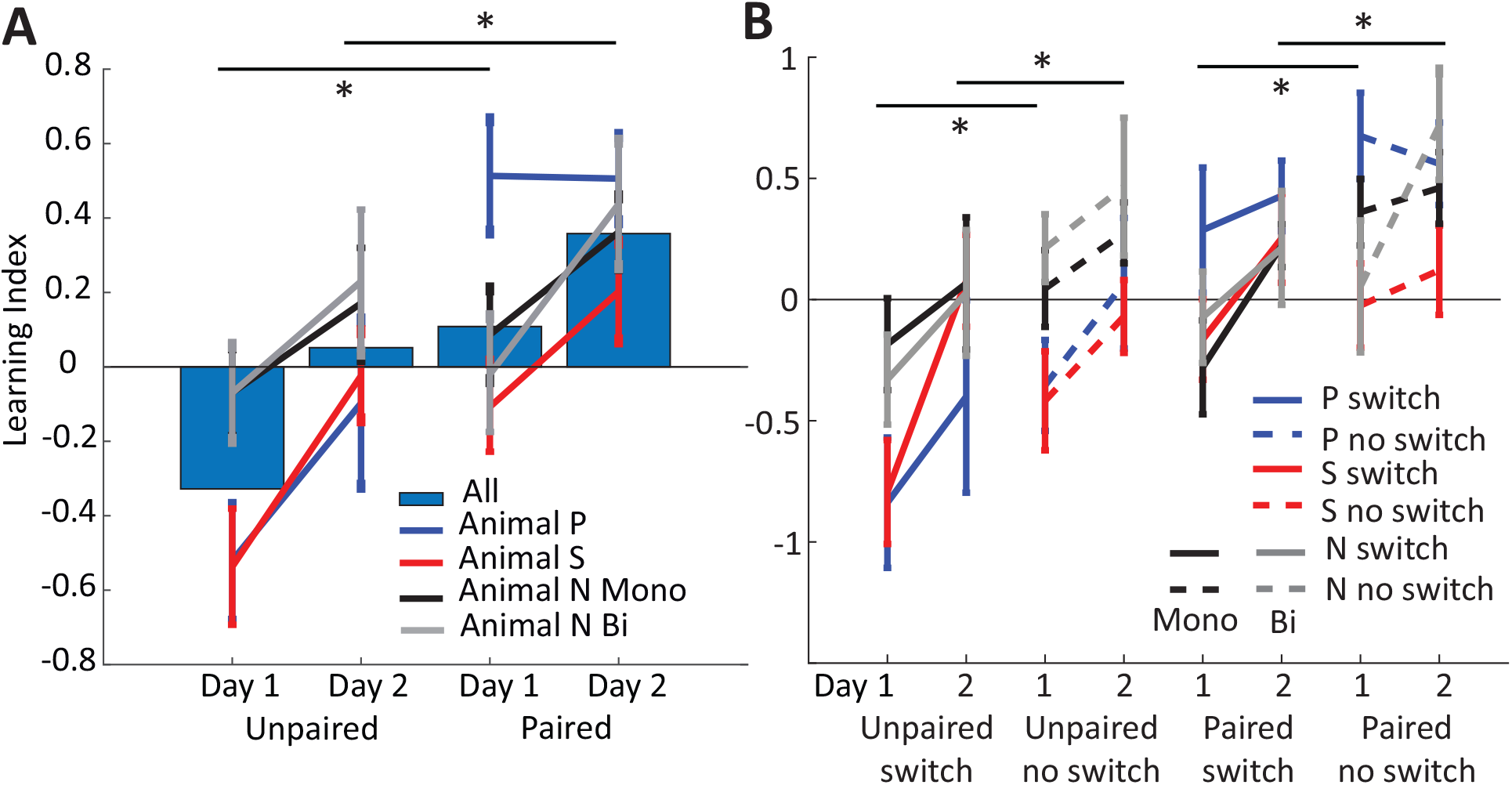
Mean learning index (LI) was larger for paired VNS-tone sessions compared to unpaired VNS-tone sessions. (A) Mean LI for each day and VNS pairing condition is plotted per animal (lines) and across all animals (blue bars). Animals often learned to respond preferentially (*LI >* 0) to the rewarded target by day 2, but LI was significantly greater when VNS was paired with tone presentation. The learning speeds (slope between day 1 and day 2) were comparable for both paired and unpaired VNS-tone conditions. Asterisks (*) indicate significant differences between means (p < 0.05, rmANOVA). (B) When LI was further subdivided into switch (more difficult, relative T1/T2 frequency reversed from previous session) and no switch (less difficult, relative frequency not reversed) conditions, overall LI was different, but learning speeds between day 1 and 2 were similar for both conditions. No significant interaction between task difficulty and day or VNS was observed (rmANOVA, see text for details). Vertical lines indicate standard error of the mean.

To summarize learning rates, we averaged LI for all paired versus unpaired VNS-target conditions, separately for day 1 versus 2 and for each animal (Fig. 6A). For all three animals, LI was higher for the paired VNS-target condition compared to that of the unpaired condition (F = 25.79, p ≈ 0, rmANOVA). LI was also consistently higher on day 1 versus day 2 for both conditions (F = 18.51, p ≈ 0, rmANOVA). However, there was no interaction between VNS condition and training day (F = 0.79, p = 0.36, rmANOVA). One of the ferrets, animal N, was trained with either monopolar (only one lead connected to the electrode cuff while the other was grounded to the body) or bipolar (both leads connected to the electrode) VNS stimulation. Overall, the results of monopolar or bipolar VNS stimulation were similar.

We also measured effects of task difficulty on LI, *i.e.*, comparing between conditions in which the relative frequency of T1 and T2 was switched between training sessions or not switched (Fig. 6B). On average, LI was lower for the more difficult switch condition in both paired and unpaired VNS conditions and on both day 1 and day 2 (F = 11.39, p = 0.0008, rmANOVA). The effect of task difficulty did not significantly interact with either VNS condition (F = 0.24, p = 0.62, rmANOVA) or training day (F = 2.7, p = 0.15, rmANOVA). However, the marginal effects of VNS condition (F = 27.4, p ≈ 0, rmANOVA) and training day (F = 19.27, p ≈ 0, rmANOVA) both remained significant.

### 3.3 Reduced responses in A1 neurons following VNS paired with BF tone

To determine effects of VNS on auditory coding, we recorded spiking activity from A1 of the three animals (P, S and N) before (pre-VNS) and after (post-VNS) during passive listening. Across a total of 201 single- and multi-units in A1, 110 had best frequency (BF) that overlapped with one of the presented tones. We tested three tone-VNS pairing conditions: (1) VNS paired with BF tone, (2) VNS paired with off-BF tone and (3) VNS during intervals between BF tone presentations. Spiking activity was not recorded during VNS due to the presence of stimulation artifacts and the possibly of current leaking into the recording system. The tone stimuli and VNS current and timing were matched to those used during behavior. We compared PSTH responses to tones pre- and post-VNS. In each non-VNS session (*e.g.*, pre- or post-VNS), there were total of 20 trials of BF tone and off-BF tone. For VNS sessions, a single tone was presented 20 times, with synchronous or asynchronous VNS, as described above. Fig. 7 shows the PSTH response of a neuron to the BF tone before and after VNS paired with the BF tone (condition 1, above).

**Figure 7.**
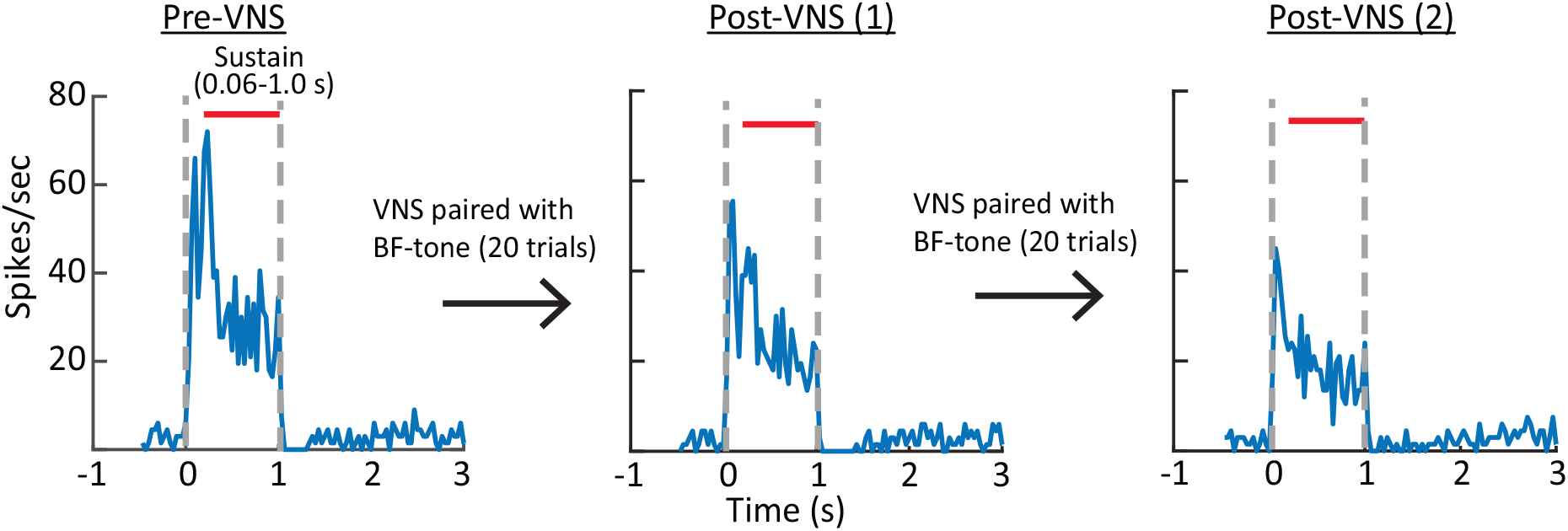
Peri-stimulus time histogram (PSTH) response of an example neuron to a tone at its best frequency (BF) before pairing (left), after one 20-trial session pairing VNS with the BF tone (middle), and after a second 20-trial pairing (right). The PSTH response was reduced following each pairing session.

Since A1 excitability is correlated with the pupil-indexed arousal state (Schwartz et al., 2019) and VNS can increase pupil size by itself (Fig. 3), we considered two possible pathways by which VNS might mediate changes in A1 responses. One pathway is coupled with pupil-indexed arousal state and induces reversible changes in A1 excitability (Fig. 1). The second pathway promotes plasticity in A1 that persists even after pupil-indexed arousal returns to its original level.

To study average changes in pupil diameter across experiments, we normalized pupil by the mean diameter on the first 5 trials of the pre-VNS session. On average, pupil diameter of each session decreased as the number of trial increased (8A). The large increase in pupil size at the beginning of post-VNS session probably reflected increased arousal after the experimenter entered the anechoic chamber to unplug the stimulation system. VNS-induced changes in pupil were smaller and shorter in duration (see Fig. 3). To isolate effects of VNS, therefore, it is important to separate changes in A1 activity into two components, one that could be predicted by fluctuations in pupil (pupil effects) and one that persisted after VNS and was not accounted for by pupil (persistent VNS effects). When we plotted the mean response of pupil and VNS effects for significant units in VNS paired with BF tone sessions, respectively, from trial to trial (Fig. 8B and C), a larger decrease in mean response of VNS effects was observed throughout the whole post-VNS session. On the other hand, the plot of mean response for each trial due to the estimated pupil effects was comparable between pre- and post-VNS sessions.

**Figure 8.**
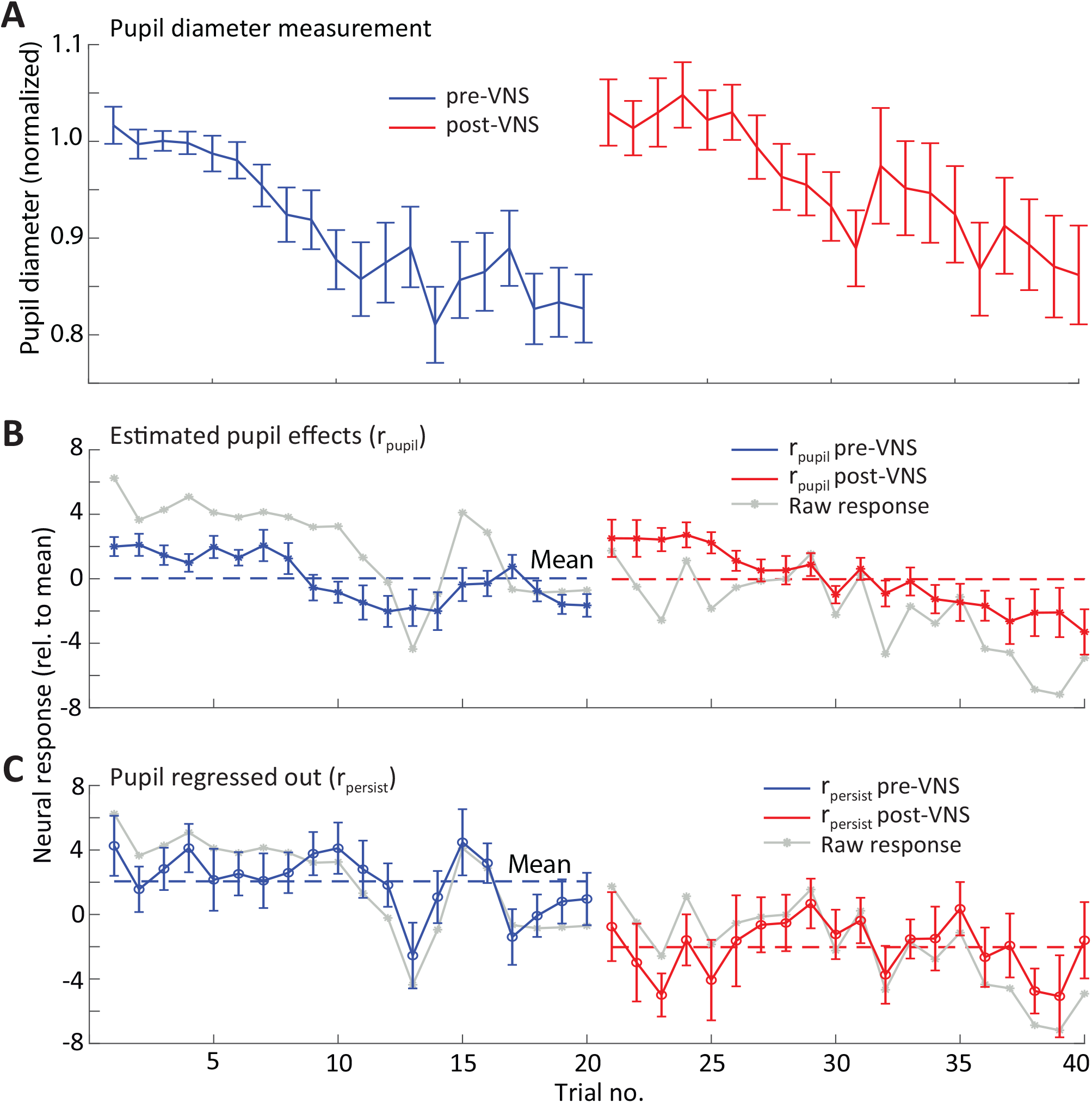
A larger decrease in mean response of VNS effects was observed during the post-VNS session. (A) Pupil diameter was normalized to the mean diameter of the first 5 trials during the pre-VNS session and then averaged across neurons (*n* = 34 neurons with significant response changes after VNS). Normalized pupil diameter of pre- and post-VNS sessions decreased as trial number increased. The mean change in BF response on each trial was separated into (B) the component that could be explained by pupil and (C) the persistent effect that could not be explained by pupil (*n* = 34). The latter component reflects a persistent change in response following VNS. Mean change in the raw response (gray line) is overlaid for comparison. The dashed blue and red lines represent the mean neural response pre- and post-VNS, respectively. Vertical lines indicate standard error of the mean.

We used a linear regression model to dissociate the effects of pupil (*r*_*pupil*_) and VNS (*r*_*persist*_) on A1 responses to the BF tone. Spike rate in *r*_*pupil*_ was predicted as the average sustained response across all trials, scaled by pupil diameter on each trial (Eq. 2). After subtracting out *r*_*pupil*_, the pre-/post- difference in the residual response, *r*_*persist*_ reflected persistent changes following VNS that could not be explained by changes in pupil.

We compared the change in the raw sustained response for each neuron following VNS-BF pairing (Fig. 9A). Across the entire set of A1 neurons, 34/110 showed a significant change post-VNS (*p* < 0.05, rank sum test) and 11/110 had mean sustained responses not were not significantly larger than 0. Across the 34 neurons with a difference, the sustained response was significantly reduced when VNS paired with BF tone (*p* = 0.035, sign rank test). After regressing out pupil a smaller number of neurons showed significant changes (23/110, Fig 9B), but the change post-VNS remained significantly negative (*p* = 0.006, sign rank test, Fig 9D). In contrast, the mean residual sustained response was not significantly reduced or increased when VNS was paired with an off-BF tone (Fig 9C). There was also not a significant change in the mean response for the unpaired condition, when VNS occurred during the inter-trial interval (results summarized in Fig. 9E).

**Figure 9.**
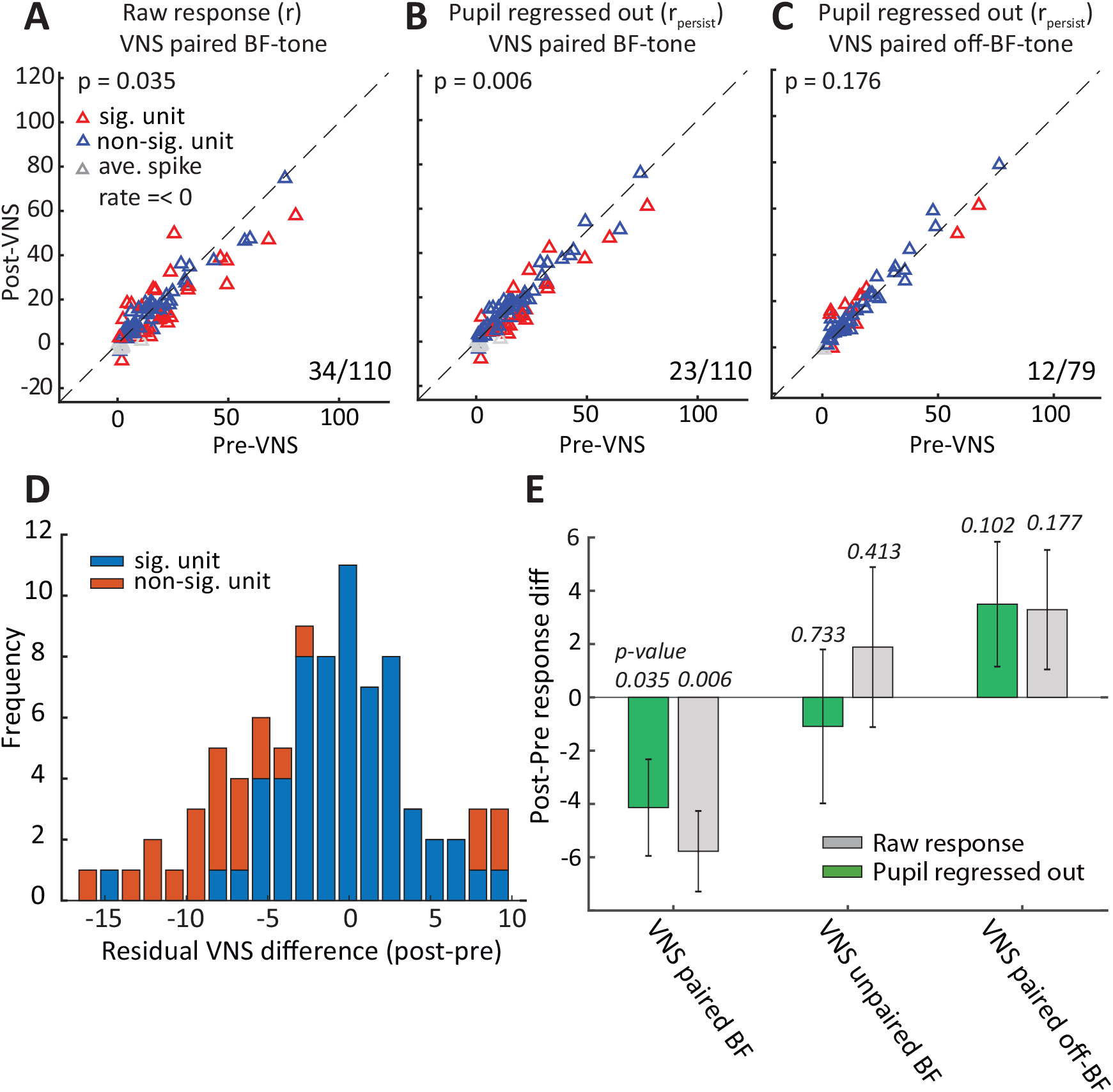
Reduced sustained response in A1 after pairing VNS with BF tone presentation. (A) Scatter plot compares the the sustained response to the BF for each A1 unit before and after pairing VNS with the BF tone. Red markers indicate units with a significant difference (*p <* 0.05, rank sum test). Numbers (right lower corner) indicate counts of neurons with significant difference. (B) Scatter plot comparing difference in sustained response after regressing out changes that can be explained by fluctuations in pupil-indexed arousal, plotted as in the left panel. (C) Scatter plot of sustained response before and after pairing VNS with an off-BF tone. As in the middle panel, effects of pupil size on spike rate were removed by linear regression. (D) Histogram of the difference in spike rate after pupil correction shows most of the units (red) with significant changes fell on the left side of the histogram. (E) Mean difference in sustained response post- versus pre-VNS, for units with significant changes in auditory responses, under different VNS pairing conditions. Numbers above each bar indicate significance of the mean change (sign rank test). Vertical lines on bars indicate standard error of the mean.

For a subset of recordings from animal N, we repeated two sessions of 20 trials pairing VNS with BF tone. Units with significant changes after either 20 or 40 trials of VNS (*p <* 0.05, Bonferoni-corrected ranksum test) were selected for pre* versus post-VNS comparison. The plot in Fig. 10A compares pupil-corrected changes in sustained response to the BF tone. For this small subset, the decrease in response after VNS was not significant. Nonetheless, the mean response showed a trend toward greater reduction as the number of VNS trials increased.

**Figure 10.**
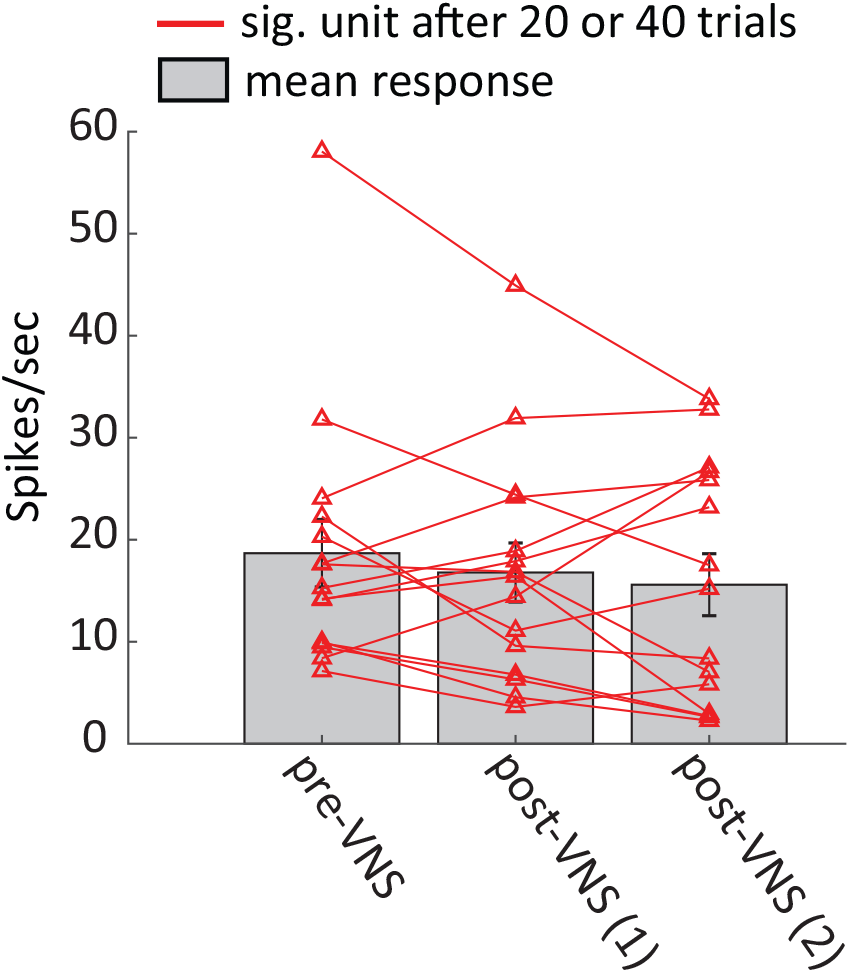
Reduced mean response for neurons with significant changes (*p <* 0.05, Bonferoni-corrected ranksum test) after first or second session of 20 trials pairing VNS with BF tone. Most neurons with significant changes show a trend toward decreased residual (pupil-corrected) response as the number of paired VNS-tone trials increased from 20 to 40. Vertical lines on bars indicate standard error of the mean.

### 3.4 Pupil changes during VNS predict persistent changes post-VNS

The results above demonstrate a relationship between VNS and short-term changes in A1 excitability that are, in turn, correlated with changes in pupil size (Fig. 9, (Schwartz et al., 2019)). Changes in pupil are associated with neuromodulatory activity (noradrenaline and acetylcholine, (Reimer et al., 2016)) which in turn is known to mediate cortical plasticity (Engineer et al., 2013; Dorr and Debonnel, 2006). Thus, we considered the possibility that persistent changes in spiking activity following VNS could be be predicted by changes in pupil size during VNS. Compared to sessions without VNS, a larger increase in pupil size was observed during the session of VNS paired with BF tone (Fig. 11A), consistent with the VNS-evoked dilation reported above (Fig. 3). Moreover, pupil dilation was especially larger during recordings when units underwent significant persistent changes post-VNS (compare red and blue lines in Fig. 11A). When we compared the mean trial evoked dilation during VNS-BF tone sessions and the magnitude of the persistent change in BF tone response, there was a significant positive correlation (*R* = 0.32, *p* = 0.006; Fig. 11B). These results indicate that changes in pupil size during VNS predicted the magnitude of persistent plasticity following VNS. This result is consistent with the possibility that changes neuromodulatory tone reflected in the pupil gate the long-term effects of VNS on sound-evoked activity.

**Figure 11.**
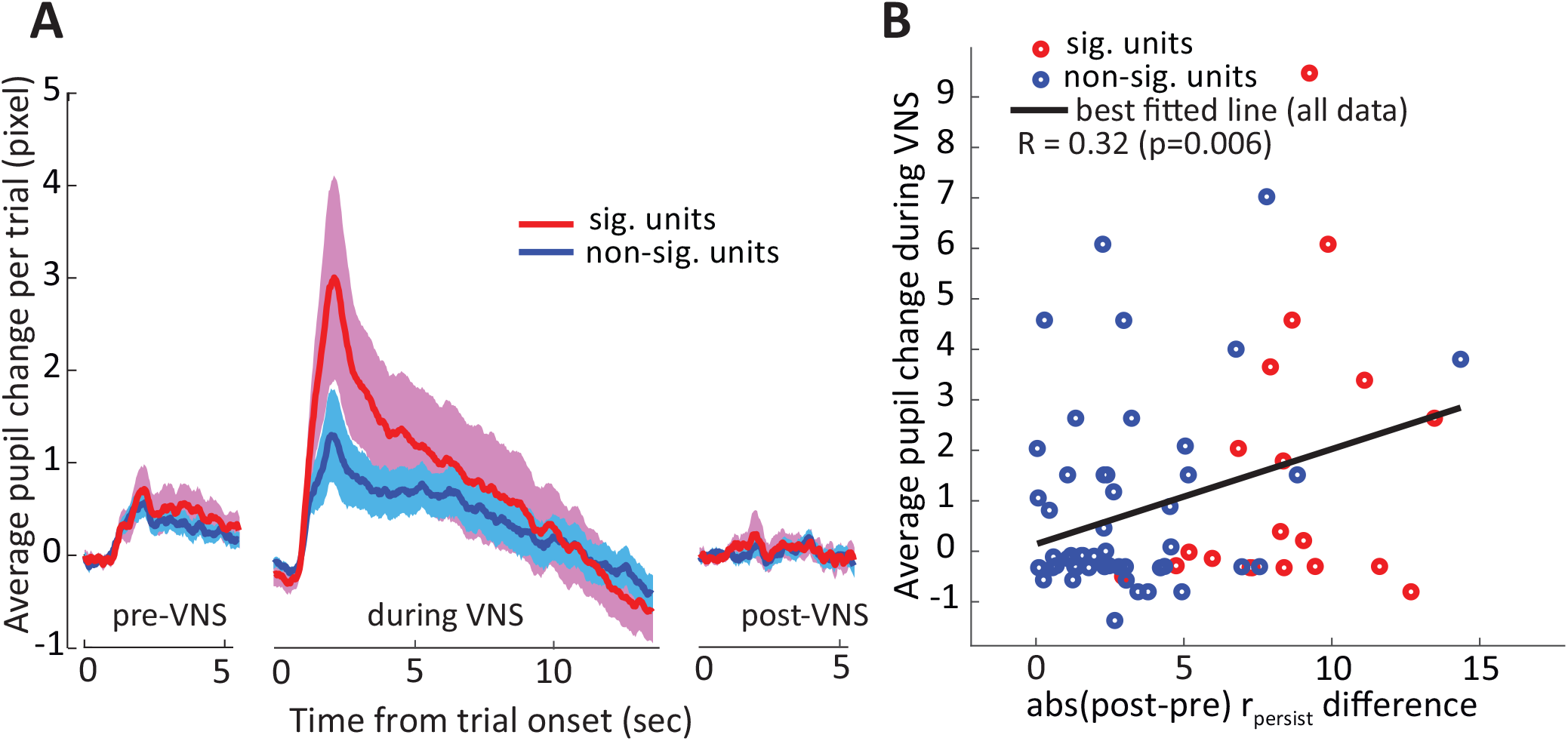
Units with significant changes between pre- and post-VNS sessions showed larger changes in pupil size during VNS paired with BF tone session. (A) Average change in pupil diameter on each trial, after subtracting to mean pupil size during the first 0.8 s of each trial. Pupil changes were grouped by sessions in which units underwent persistent changes post-VNS (red) or did not (blue). Changes in pupil were always large during VNS sessions, but the changes were especially larger during sessions that produced persistent changes in neural responses. Shading indicates standard error of the mean. (B) Scatter plot compares the magnitude of persistent changes in neural response post-VNS against trial-evoked change in pupil during the preceding VNS (*R* = 0.32, *p* = 0.006).

## 4 DISCUSSION

Previous studies have shown that extended periods of pairing VNS with acoustic stimuli (300 times/day for 20 days) can induce plasticity in sound coding by primary auditory cortex (A1) (Buell et al., 2018; Borland et al., 2016; Engineer et al., 2011, 2015b) and auditory midbrain (Borland et al., 2019). Building on this work, we characterized short-term effects of VNS (200 times/day for 1-2 days) on auditory learning and stimulus-specific activity in A1. We found that when VNS was paired with target tones during classical conditioning, animals took fewer trials to learn to respond selectively to the rewarded tone than in an unpaired VNS condition. Evidence for faster learning appeared consistently, across multiple animals and difficulty conditions, even on the first day of a new reward pairing condition. These results demonstrate that acute periods of VNS can have significant impact on auditory learning.

In addition to enhanced learning on short timescales, we found that A1 neurons in passive listening animals showed reduced responses after a similar pattern of VNS as during behavior. After 20 trials of pairing VNS with a BF tone, the mean sustained response to the BF tone was reduced relative to before VNS. This reduction in response persisted even after regressing out changes in spiking that could be explained by fluctuations in the pupil that index changes in global arousal. Moreover, the size of pupil fluctuations during VNS correlated with the magnitude of the persistent change in neural response following VNS. These results support the idea that the rapid and reversible effects of VNS, which are indexed by pupil diameter, positively gate long-term plasticity in A1 (see Fig. 1). These results provide new insight into short-term effects of VNS and tease apart the pathways that mediate its effects on auditory processing.

### 4.1 Impact of VNS on behavioral training and learning

Although research into the effects of VNS on auditory learning is limited (Mridha et al., 2019), there is substantial evidence for positive effects of pairing VNS with rehabilitation during recovery from motor disorders (Khodaparast et al., 2016, 2014; Hays et al., 2014a,b). In a study conducted by Khodaparast *et al.* (2014), brief VNS was delivered with each successful movement (within 0.5 sec) during rehabilitation in rats with cortical ischemia, and a complete recovery of forelimb function to pre-lesion levels was observed (Khodaparast et al., 2014). In contrast, animals receiving VNS after rehabilitation or receiving rehabilitative training alone failed to restore function to pre-lesion levels. VNS has also been demonstrated to reduce conditioned fear when paired with conditioned cue during fear extinction in rats (Pena et al., 2013, 2014). Even when extinction training was delayed for 2 weeks after initial fear conditioning, animals receiving paired VNS showed significantly less fear response than sham control rats after a single day of extinction training (Pena et al., 2013). This finding is consistent with our results showing animals receiving paired VNS-tone during training responded more to the rewarded tone and showed significant learning effects after a single day of training (Fig. 6).

Previous studies of VNS in auditory learning have involved relatively simple acoustic discrimination or detection (Pena et al., 2013; Mridha et al., 2019). Similarly, in the current study, rewarded and non-rewarded tones were separated by > 1 octave. Thus the major impact of VNS may have been on learning stimulus-reward associations rather than the discrimination of very similar stimuli. Pena *et al.* (2014) provide evidence that VNS enhances extinction not only for the stimulus paired with VNS but also for another conditioned stimulus associated with the same fear experience but not paired with VNS (Pena et al., 2014). Thus, it remains unclear whether VNS-sound pairing is more beneficial to learning new acoustic categories or learning reward associations with known categories.

### 4.2 Importance of VNS timing and neuromodulator release in plasticity

Stimulation of vagus nerve triggers the release of neuromodulators from multiple nuclei to drive plasticity, including noradrenergic (LC), cholinergic (NB) and serotonergic (dorsal raphe nucleus) systems (Nichols et al., 2011; Dorr and Debonnel, 2006; Hulsey et al., 2019). A reduction of either noradrenergic or cholinergic signaling prevents VNS-dependent effects in the central nervous system, further suggesting that VNS engages these systems (Nichols et al., 2011; Krahl et al., 1998). Furthermore, the importance of precise timing of VNS in the induction of plasticity has also been emphasized in many studies (Engineer et al., 2011; Porter et al., 2012). Neuroplasticity is strongly influenced by the relative timing of stimuli and neuromodulator release. As a result, pairing VNS with a sensory input, for example a movement or an acoustic stimuli, improves recovery from brain injury or tinnitus by enhancing neuroplasticity in a timing-dependent manner (Hays et al., 2013; Pruitt et al., 2016; Khodaparast et al., 2016, 2014). In our results, we found that animals learned reward associations faster in the paired VNS-target condition when VNS was synchronized with target tone presentation. This finding is consistent with the hypothesis that precise timing of neuromodulator release with respect to behaviorally-relevant sensory stimulation is required for the beneficial neuroplasticity associated with auditory learning.

### 4.3 VNS effects on primary auditory cortex

There is growing evidence of parallels between effects of VNS and of direct neuromodulatory stimulation. In their seminal work on auditory plasticity, Kilgard *et al.* (2002) demonstrated that electrical stimulation of the cholinergic basal forebrain (nucleus basalis, NB) while simultaneously delivering a sensory stimulus drives plasticity in auditory cortex that mimics plasticity induced by perceptual learning Kilgard et al. (2002). Exposing a rat to a fixed-frequency tone paired with chronic stimulation of the NB (300 time/day for 20 days) resulted in large-scale map reorganization of the auditory cortex that was specific to frequencies near the paired tone (Kilgard and Merzenich, 1998). Furthermore, by pairing more complex stimuli with NB stimulation, selective enhancement has been produced in A1 for a wide range of acoustic features, including sound level, temporal modulation, and frequency (Kilgard and Merzenich, 1998; Kilgard et al., 2002,?, 2001; Pandya et al., 2005; Moucha and Kilgard, 2006).

Since afferent fibers of the vagus nerve innervate cells in the nucleus tractus solitarius, which in turn projects to LC and NB (see Fig. 1 in Engineer *et al.*, 2012), electrical stimulation of the vagus nerve should have effects similar to direct stimulation of the NB (Engineer et al., 2011, 2013). Indeed, pairing VNS with a 9 kHz tone caused a 79 % increase in the number of A1 neurons with a characteristic frequency near the paired tone compared to naive control rats (Engineer et al., 2013). Moreover, repeatedly pairing VNS with rapid 15 pps tone trains increased the temporal following rate of A1 neurons while pairing VNS with slow 5 pps tone trains decreased the temporal following rate (Engineer et al., 2013; Shetake et al., 2012). These effects can generalize across stimuli. After pairing tone trains at rapid 15 pps with VNS, A1 responses were also increased for unpaired novel speech sounds (Engineer et al., 2017). It has also been demonstrated that VNS modulates synchrony and excitability in the A1 at least in part through the activation of muscarinic acetylcholine receptors (Nichols et al., 2011).

The general observation from studies that perform chronic VNS has been of enhanced firing rate of stimulus-specific neural A1 responses (Engineer et al., 2015b; Borland et al., 2019). However, the short-term effects of VNS may be different. Nichols *et al.* (2011) reported that shorter bouts of VNS increased and decorrelated spontaneous activity of A1 neurons and suppressed entrainment to repeated 6-8 Hz noise stimulation. This study recorded multi-unit activity in layers 4/5 of anesthetized rat A1 and performed 100 repetitions of VNS (500 ms train of 500 *μ*s biphasic pulses at 30 Hz) repeated every 10 sec (Nichols et al., 2011). In the current study, we recorded both single- and multi-unit activity in A1 of awake animals. We performed 20 or 40 repetitions of VNS (1 sec train of 200 *μ*s biphasic pulses at 30 Hz) repeated every 12.5 sec. Despite some differences in methodology and parameters used for acute VNS, we also observed a suppression of evoked A1 responses after VNS (Figs. 7 and 9, similar to the observation by Nichols *et al.* (2011).

The explanation for why short-term VNS leads to suppressed A1 responses while long-term VNS tends to enhance responses remains unclear. In addition to opposite effects on evoked activity, changes in spontaneous rate also differ. According to Borland *et al.* (2019), spontaneous rate was reduced in A1 of rats after long-term VNS-tone pairing, contrasting with the increase reported by Nichols *et al.* (2011). These differences could reflect a non-monotonic relationship between the number of VNS trials and subsequent plasticity. Alternatively, slow compensatory processes, such as changes in inhibitory network tone, could alter evoked activity over a longer time following VNS. Also possibly relevant, many recordings following chronic VNS have been performed in anesthetized animals, and changes in A1 responses induced by VNS could be masked or affected by anesthesia (Cheung et al., 2001).

### 4.4 Pupil dilation and gating of long-term VNS effects

Recent work has shown that neuromodulatory activity is correlated with luminance-independent changes in pupil diameter (Joshi et al., 2016; Murphy et al., 2014; Desbeaumes Jodoin et al., 2015). Spontaneous fluctuations in pupil size are correlated with changes in sensory cortical activity (McGinley et al., 2015; Vinck et al., 2015) and track rapid changes in activity of adrenergic and cholinergic axon terminals in cortex (Reimer et al., 2016). Pupil dilation has also been observed following VNS (Bianca and Komisaruk, 2007; Desbeaumes Jodoin et al., 2015) and has been proposed as a biomarker for effective stimulation (Mridha et al., 2019). Because both VNS and changes in pupil diameter are associated with fluctuations in cortical activity, we considered the possibility that acute effects of VNS in A1 could be explained by the global changes in arousal registered in pupil dilation. To control for this possibility, we regressed out the pupil effects using linear regression. We observed significant residual changes in A1 activity (Fig. 9), even after the regression.

Moreover, when we analyzed the correlation between pupil changes during VNS and the persistent changes in A1 activity following VNS, we found a significant correlation (Fig. 11). Given the association we observe between the size of VNS-evoked changes in pupil and persistent changes in A1 responses, we speculate that plasticity in A1 is mediated by two pathways, namely a short-term, reversible pathway and a long-term, persistent pathway (Fig. 1). In the short-term pathway, changes in pupil correlate with A1 excitability (Schwartz et al., 2019). In the long-term pathway, which maybe gated by the short-term pathway, VNS promotes persistent A1 plasticity that is related to learning improvement. Both short- and long-term effects can be driven by VNS-mediated changes in the occur downstream of VNS via changes in neuromodulation.

## 5 CONCLUSION

We find that acute stimulation of the afferent vagus nerve facilitates learning of sound-reward associations and modulates stimulus-specific activity in A1. In a simple classical conditioning paradigm, the rate with which ferrets learned to anticipate rewards following one of two tones increased when the tones were paired with VNS. The benefits of VNS were evident even on the first day following introduction of a new tone-reward association. In addition, neurophysiological recording in A1 of passively listening animals revealed a decrease in sustained responses to BF tones after acute stimulation when VNS was paired with that tone. VNS produced rapid, reversible changes in pupil diameter, consistent with a general increase in arousal. The size of these changes in pupil diameter predicted the magnitude of persistent changes in sound-evoked activity following VNS, suggesting that the rapid, reversible changes in neuromodulation associated with pupil dilation may gate long-term and persistent plasticity induced by VNS in auditory cortex. Appropriately timed VNS during learning may increase the speed with which adults learn to use auditory prosthetics or acquire new languages.

## CONFLICT OF INTEREST STATEMENT

The authors declare that the research was conducted in the absence of any commercial or financial relationships that could be construed as a potential conflict of interest.

## AUTHOR CONTRIBUTIONS

Jesyin Lai and Stephen V. David designed research; Jesyin Lai performed research; Jesyin Lai analyzed data; Jesyin Lai and Stephen V. David wrote and edited the paper.

## FUNDING

This work was supported by grants from DARPA (D15AP00101) and National Institution Deafness and Other Communication Disorders (NIDCD R01 DC014950).

## ACKNOWLEDGMENTS

The authors would like to thank Luke A. Shaheen in assistance with cuff electrode testing and data analysis of array recording; Luke A. Shaheen, Zachary P. Schwartz and Daniela Saderi for training and assistance with neurophysiological recording; Zachary P. Schwartz and Charles R. Heller for assistance with pupillometry.

